# Identification of phosphosites that alter protein thermal stability

**DOI:** 10.1101/2020.01.14.904300

**Authors:** Ian R. Smith, Kyle N. Hess, Anna A. Bakhtina, Anthony S. Valente, Ricard A. Rodríguez-Mias, Judit Villén

## Abstract

Proteomics has enabled the cataloguing of 100,000s of protein phosphorylation sites ^1^, however we lack methods to systematically annotate their function. Phosphorylation has numerous biological functions, yet biochemically all involve changes in protein structure and interactions. These biochemical changes can be recapitulated by measuring the difference in stability between the protein and the phosphoprotein. Building on recent work, we present a method to infer phosphosite functionality by reliably measuring such differences at the proteomic scale.

## MAIN TEXT

Recently, Huang et al.^2^ developed the Hotspot Thermal Profiling (HTP) method to identify phosphosites that alter protein thermal stability, reporting 719 out of 2,883 (25%) phosphosites with significant effects. The reported melting temperatures (T_m_) for phosphopeptides correlated poorly with the T_m_ for their corresponding proteins (R^2^ = 0.18) (Fig. 1a), implying that many phosphosites function by structurally reshaping the proteome. However, the low T_m_ reproducibility between replicates (Supplementary Fig. 1a) suggests that this conclusion may be due to technical variation (Supplementary Discussion). The HTP workflow consists of phosphopeptide enrichment followed by separate isotopic labeling and mass spectrometric analysis to derive T_m_ values for phosphopeptides and proteins, respectively. Because phosphopeptide samples also contained unmodified peptides, which are expected to have the same T_m_ as the protein, we can use these peptides to assess technical variation between the two samples. Disconcertingly, our re-analysis revealed that 626 out of 3074 (20%) of the co-enriched unmodified peptides had significant stability effects, almost the same percentage as phosphopeptides (22%, 596 out of 2656 in our re-analysis) (Fig. 1b, Dataset S1). Additionally, the T_m_ correlation of these peptides with their protein T_m_ was similarly low (R^2^ = 0.18) to the correlation between phosphopeptides and protein (Supplementary Fig. 1b). In the absence of a biological explanation, this suggests that the independent labeling and mass spectrometric analysis of peptide and phosphopeptide samples could have introduced substantial technical error precluding the comparison, and perhaps that the reported hits arise from a lack of stringency in the applied statistical analysis (Supplementary Discussion).

**Figure 1:**
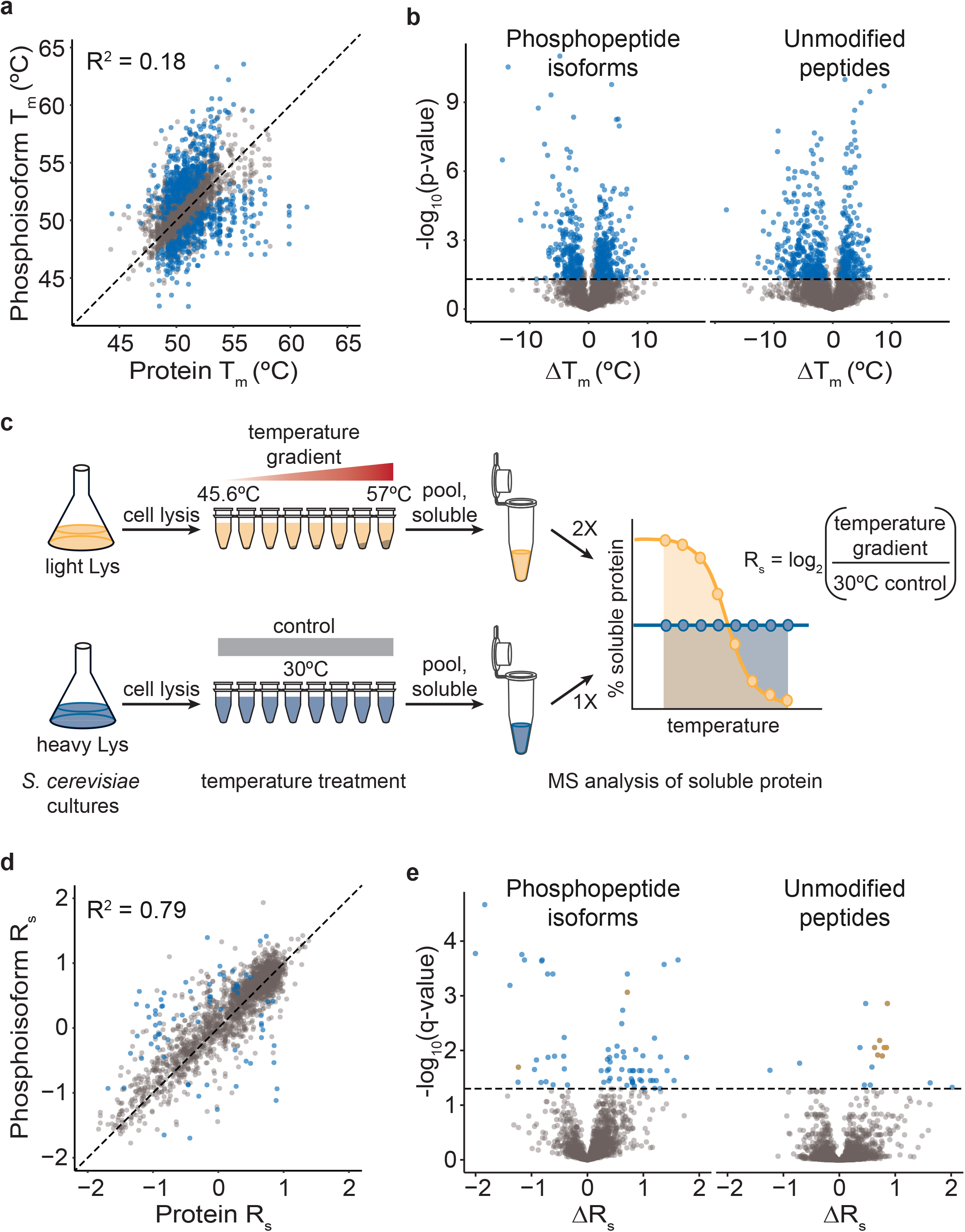
Most phosphosites have little effect on protein stability. **a**, Scatter plot and Pearson correlation between T_m_ of phosphopeptide isoforms (n=10) and T_m_ of the corresponding protein (n=11). Values were obtained from the Huang et al. supplementary dataset, using mean T_m_ values. Significant phosphopeptide isoforms in blue. **b**, Volcano plots showing differences in protein thermal stability (T_m_) between phosphopeptide isoforms (n=10, left panel) or unmodified peptides observed in the phosphopeptide enriched sample (n=10, right panel) and their corresponding protein (n=10) (t-test, significant hits at p-value <0.05 in blue), from Huang et al. data reanalysis. **c**, Dali workflow depicting SILAC labeling of yeast, the gradient temperature treatment of the protein extract, the inclusion of a 30°C control, the quantification of soluble protein by mass spectrometry, and the calculation of relative stability (R_s_). **d**, Scatter plot and Pearson correlation as in (b) using Dali’s R_s_ data. **e**, Volcano plots as in (a) with x-axis showing R_s_ values. Here n=6, p-values were Benjamini-Hochberg corrected and significant hits at q-value < 0.05. Significant hits in blue, significant phosphopeptide isoforms found in proteins with known cleavage or splicing events are in orange.

To minimize technical noise derived from sample preparation, peptide samples should be labeled and mixed prior to phosphopeptide enrichment (Supplementary Discussion and accompanying manuscript^3^). Because scaling-up isobaric chemical labeling increases reagent costs substantially, we have developed an alternative approach to identify phosphosites that alter thermal stability, that we call Dali (Fig. 1c). Dali applies the Proteome Integral Stability Alteration (PISA) method^4^, a simplified version of thermal proteome profiling^5^, in which the soluble protein from the different temperature points are combined to provide an estimation of the area under the protein melting curve. To reliably compare phosphopeptides to proteins, we normalize each measurement to a 30°C treated proteome reference that is labeled with heavy lysine, obtaining a relative stability (R_s_) measurement for phosphopeptides and proteins. This 30°C reference is mixed in with the temperature gradient treated samples prior to protein digestion, and it is present during phosphopeptide enrichment and mass spectrometry (MS) measurement of peptides and phosphopeptides.

We applied Dali to the *S. cerevisiae* proteome and obtained reproducible R_s_ measurements for proteins (average R^2^= 0.76) and phosphopeptides (average R^2^= 0.65) (Supplementary Fig. 1a). In contrast to the Huang et al. dataset, we find that the stability of phosphopeptides correlates well with the stability of their respective proteins (R^2^=0.79 for mean R_s_ comparisons) (Fig. 1d), suggesting that most phosphosites do not alter protein stability as also observed by Potel et al.^3^. As expected, the stability of non-modified peptides present in the phosphopeptide-enriched samples also correlated well with their proteins (R^2^ = 0.90 for mean R_s_ comparisons), indicating that R_s_ measurements in the phosphopeptide samples and protein samples can be reliably compared (Supplementary Fig. 1c). Finally, our analysis yielded 71 phosphopeptide isoforms out of 2,345 (3%) with significantly different thermal stability than the unmodified protein (Fig. 1e, Dataset S2). We detected several non-modified peptides with significant differences in stability, yet this set constituted a much smaller fraction than found in Huang et al. (Dataset S3). Many of these peptides (7 out of 16) were cases where the protein is known to be post-translationally processed via cleavage (e.g. RPS31^6^) or splicing (e.g. VMA1^7^), resulting in proteins and/or proteoforms of different thermal stability as our method measured (Supplementary Fig. 2).

Among phosphosites that decreased protein thermal stability, we identified four sites located at protein interfaces (Ser56 on PUP2, Ser59 on ARO8, Ser79 on TPI1 and Ser201 on GAPDH) (Fig. 2a, Supplementary Fig. 3) that may act by disrupting protein-protein interactions. For example, PUP2 is the alpha 5 subunit of the 20S proteasome, and Ser56 is a known Cdc28 substrate^8^ located at the protein interaction interface with PRE6, the 20S proteasome alpha 4 subunit (Fig. 2a). The stability measured for the phosphopeptide spanning Ser56 is significantly lower than the stability of PUP2, which is similar to other proteins in the 20S proteasome, suggesting Ser56 phosphorylation may dissociate PUP2 from the 20S proteasome.

**Figure 2:**
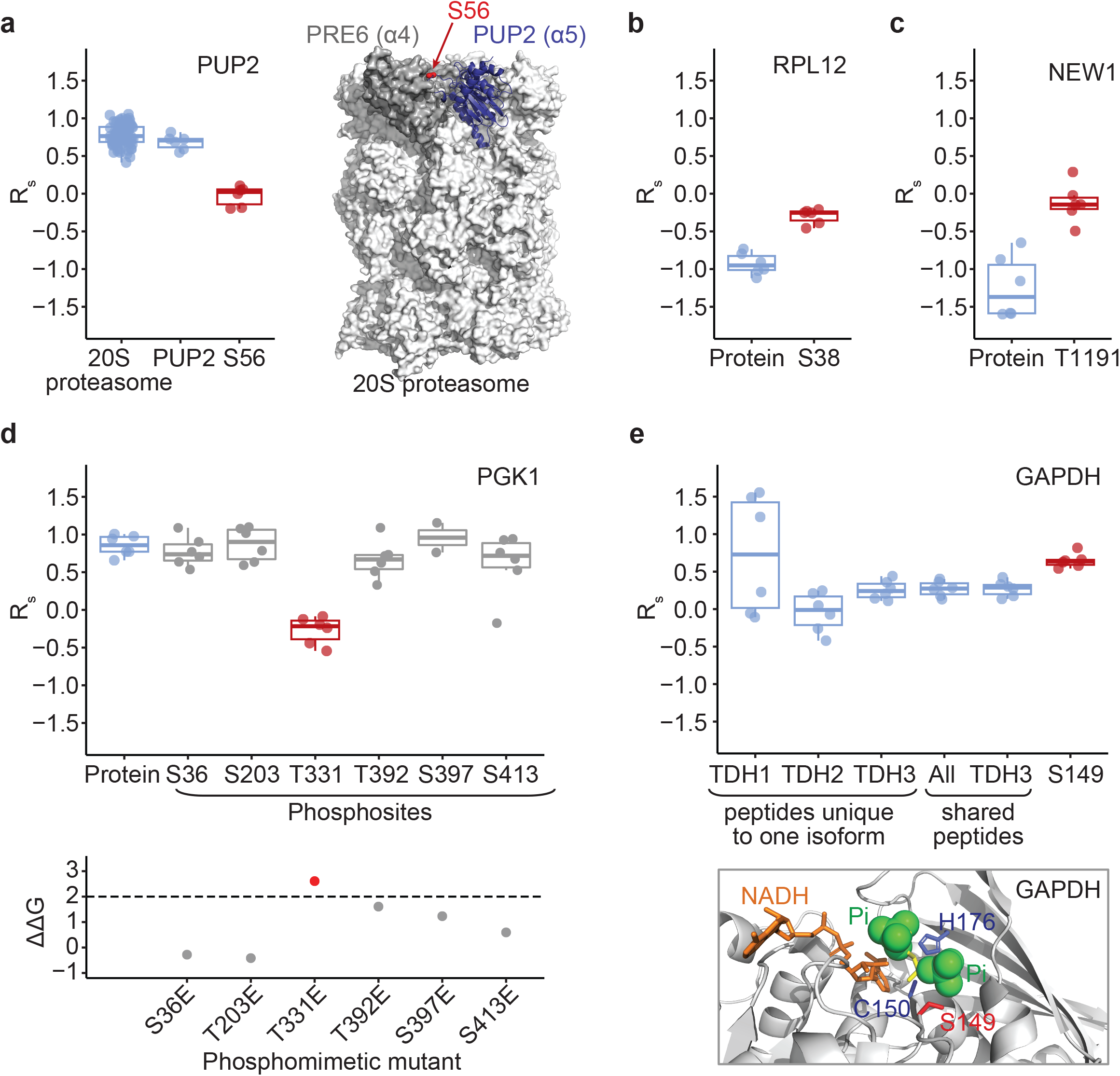
Examples of phosphosites that alter protein thermal stability. **a**, Boxplot of R_s_ values and distributions for the phosphopeptide containing PUP2 Ser56, PUP2 protein, and all the proteins in the 20S proteasome. Shown at the right is the structure of the 20S proteasome with PUP2 in blue, Ser56 in red, and PRE6 in grey. **b**, R_s_ values for RPL12 and RPL12 S38 phosphopeptide isoform. **c**, R_s_ values for NEW1 and phosphopeptide isoform containing NEW1 T1191. **d**, R_s_ values for PGK1 and all measured PGK1 phosphopeptide isoforms, with significantly destabilizing phosphosite S331 shown in red. ΔΔG_pred_ for all glutamic acid phosphomimetic substitutions were obtained from mutfunc, with ΔΔG_pred_ > 2 considered likely destabilizing. **e**, GAPDH S149 phosphopeptide is shared across all GAPDH paralogs (TDH1, TDH2, and TDH3). Boxplot shows R_s_ values and distributions for peptides unique to one isoform (TDH1, TDH2, TDH3), peptides shared among all GAPDH isoforms (all), all peptides for TDH3, and the S149 phosphopeptide isoform. Bottom panel shows localization of S149 on GAPDH structure near the binding site of the enzyme substrate. All boxplots show results from 6 biological replicates.

We identified stabilizing phosphosites that may play a role in the protein translation process. For example, we found that phosphorylation at Ser38 on ribosomal protein RPL12/uL11 significantly increased protein stability (Fig. 2b). This phosphosite is an evolutionarily-conserved Cdc28 substrate^8^ that is regulated during the cell cycle^9^ and has been reported to be depleted in polysomes and influence mitotic translation^10^. Due to RPL12 location at the ribosome P-stalk and the proximity of residue Ser38 to elongation factor 2, we hypothesize that this phosphosite may modulate the interaction with EF-2 to aid ribosomal translocation during protein synthesis, and the change in conformation and binding may stabilize RPL12. We also identified a stabilizing phosphorylation on NEW1 at Thr1191 (delta R_s_ = 1.23) (Fig. 2c). NEW1 is a translation factor that binds to the ribosome at a position analogous to eEF3 and fine-tunes the efficiency of translation termination^11^. The identified phosphosite fits the CK2 consensus motif, is located within the acidic C-terminal sequence of NEW1, and is highly conserved. A T1191A mutant has growth defects^12^ suggesting that phosphorylation is important for NEW1 function.

We were able to measure many key glycolysis proteins identifying phosphosites that may modulate enzyme kinetics. For example, we measured the stability for six phosphosites on PGK1, of which only Thr331 showed significantly decreased stability (Fig. 2d). This observation agrees with the predicted stability effects of phosphomimetic substitutions on PGK1^13^ (Fig. 2d). We identified a stabilizing phosphosite at Ser149 in the three GAPDH isozymes TDH1, TDH2 and TDH3 (Fig 2e). Ser149 is adjacent to catalytic Cys150 and to the binding sites of substrates glyceraldehyde-3-phosphate (G3P) and inorganic phosphate. Interestingly, Ser149 phosphorylation would occupy the inorganic phosphate binding site (Fig 2e). Additionally, it has been recently reported that a TDH3 S149A mutant exhibits a growth defect with doxorubicin compared to wild-type and decreases TDH3 activity to a greater extent than a TDH3 knockout^14^. Our results raise the possibility that S149 phosphorylation may increase the stability of apo-GAPDH, the GAPDH-G3P reaction intermediate and aid phosphate transfer by enhancing product release.

In this communication, we have outlined a novel proteomic method that enables robust thermal stability comparison between proteins and phosphorylated proteoforms. Our method identified 3% phosphosites in the *S. cerevisiae* proteome that significantly changed protein melting behavior, with several examples potentially altering protein conformation and interactions. Additional experiments will be needed to precisely characterize the function of these phosphosites. One limitation of this method is that the sensitivity to detect changes in stability is lower for proteins with extreme (low or high) melting temperature, which can be circumvented by performing the experiment using different temperature gradients. Our method can be extended to other model organisms and cell culture systems, as well as to other post-translational modifications, expanding the proteomic toolkit to functionally annotate dynamic protein modifications at scale.

## METHODS

### Yeast strains

All yeast experiments were conducted on the *Saccharomyces cerevisiae* haploid strain BY4741 (MATa his3Δ1 leu2Δ0 met15Δ0 ura3Δ0), a direct descendant from FY2, which is itself a direct descendant of S288C.

### S. cerevisiae growth, stable isotope labeling, and cell harvest

Two overnight yeast cultures were grown at 30°C in synthetic complete media (SCM) containing 6.7g/L yeast nitrogen base, 2g/L of synthetic complete mix minus lysine, 2% glucose, and supplemented with either regular lysine (light culture) or ^2^H_4_-lysine (heavy culture) at 0.872 mM final concentration. These cultures were used to seed three 50mL cultures of each light and heavy at OD_600_ 0.15, which were grown at 30°C and 45mL were harvested at OD_600_ ~ 1 by centrifugation at 7,000 × g for 10min. Yeast pellets were washed by resuspension in 1.5mL ice-cold sterile water and centrifugation in 2mL screw cap tubes at 21,000 × g for 10min; and then snap-frozen in liquid nitrogen and stored at −80°C.

### Cell lysis and protein extract temperature treatment

Frozen yeast cell pellets were resuspended in 700μL of non-denaturing lysis buffer (50mM HEPES pH 7.0, 75mM NaCl) containing 0.5X protease inhibitors (Pierce) and phosphatase inhibitors (50mM β-glycerophosphate, 10mM sodium pyrophosphate, 50mM of NaF, 1mM sodium orthovanadate) on ice. Cells were lysed by bead beating with 0.5mm zirconia/silica beads for 4 cycles of 60sec of mechanical agitation followed by 90sec rest on ice. Lysates were clarified by sequential centrifugation, first at 1,200 × g for 1min to remove the beads and then at 21,000 × g for 10min at 4°C to remove cell debris. To bring all protein extracts to the same concentration, extract volumes were adjusted to 1 OD_600_ unit from a 45mL culture in 1mL.

Each cell extract was aliquoted into 2 strips of 8 PCR tubes each (1×8 for the temperature gradient and 1×8 for the 30°C) dispensing 50μL of protein extract per tube. All samples were initially equilibrated to 30°C for 5 min. Temperature gradient samples were subjected to 45.6°C, 46.8°C, 48.3°C, 50°C, 52°C, 53.6°C, 54.9°C, and 57°C, one tube to each temperature, for 5min. In parallel, controls were subjected to an additional 30°C temperature treatment for 5min. All samples were cooled down to room temperature for 10min. For each replicate, temperature gradient samples were all pooled into one tube and 30°C controls were pooled into a separate tube prior to centrifugation at 21,000 × g for 30min at 4°C. The soluble protein fractions for the temperature gradient and 30°C controls were combined 2:1, three replicates with the temperature gradient labeled heavy and the 30°C controls labeled light, and three additional replicates with the labels swapped. We generated additional controls where heavy and light 30°C controls were combined to assess potential differences in protein expression due to the different labeling. Protein concentration was measured by the BCA assay.

### Protein reduction, alkylation, LysC digestion, and desalting

Samples were diluted 2-fold with a buffer containing 8M urea, 50mM HEPES pH 8.9, 75mM NaCl, 1mM sodium orthovanadate, 50mM β-glycerophosphate, 10mM sodium pyrophosphate, 50mM NaF. Protein samples were subjected to reduction with 7.5mM dithiothreitol (DTT) for 30min at 55°C and alkylation with iodoacetamide (22.5mM) for 30min at room temperature in the dark with agitation. The alkylation reaction was quenched with an additional 7.5mM DTT at room temperature for 30min with agitation. The pH was adjusted to 8.5 with 1M Tris pH 8.9. Lysyl endopeptidase (LysC; Wako Chemicals) was added at a 1:100 enzyme to protein ratio and protein samples were incubated overnight with agitation at room temperature. LysC digestion was quenched by addition of trifluoroacetic acid (TFA) to a final concentration of 1% and pH ~2-3 and the digests were stored at −80°C.

Peptide samples were desalted by solid-phase extraction over 50mg Sep-Pak tC_18_ cartridges (Waters). Packing material was washed with 1mL methanol, 3 × 1mL 100% acetonitrile, 1mL 70% acetonitrile, 0.25% acetic acid, 1mL 40% acetonitrile, 0.5% acetic acid, and equilibrated with 3 × 1mL 0.1% TFA. Peptides were then loaded by gravity twice, washed with 3 × 1mL 0.1% TFA and 1mL 0.5% acetic acid. Peptides were eluted with 600μL of 40% acetonitrile, 0.5% acetic acid and 400μL 70% acetonitrile, 0.25% acetic acid, and aliquoted as follows: 40μg for high-pH reversed-phase fractionation, 200μg for Fe^3+^-IMAC phosphopeptide enrichment, and 10μg for preliminary LC-MS/MS analysis to assess sample quality. All samples were dried by vacuum centrifugation and stored at −80°C.

### High-pH reversed-phase fractionation

Peptides were fractionated by high-pH reversed-phase fractionation on a 200μL pipette tip packed with 4 layers of SDB-XC material (Empore). The material was washed with 50μL methanol, 50μL 80% acetonitrile, 20mM ammonium formate, and 3 × 50μL 20mM ammonium formate. Peptides (40μg) were solubilized in 40μL of 5% acetonitrile, 20 mM ammonium formate, loaded onto the SDB-XC tip, and the flow-through was collected in a mass spectrometer vial (fraction 1). Peptide fractions 2-5 were obtained by step elution with 40μL of 20mM ammonium formate in 10 %, 15%, 20%, and 80% acetonitrile and collection in mass spectrometry vials. Peptide fractions were dried by vacuum centrifugation, solubilized in 3% acetonitrile, 4% formic acid, and ~1μg of each fraction was analyzed by LC-MS/MS.

### Fe^3+^-NTA IMAC phosphopeptide enrichment

Phosphopeptide enrichment was conducted by immobilized iron cation affinity chromatography in batch mode and automated in a 96-well format on a KingFisher magnetic particle processor as we described^15^. For each sample, ~200μg peptides were solubilized in 70 μL 0.1% TFA, 80 % acetonitrile and incubated with 80 μL of a 5% slurry of magnetic Fe-NTA beads (Cube Biotech) in the same solvent for 30min. Beads were washed three times with 150μL 0.1% TFA, 80% acetonitrile and phosphopeptides were eluted with 50μL 50% acetonitrile, 0.37M ammonium hydroxide. Eluates were acidified with 30μL 10% formic acid, 75% acetonitrile and filtered over two-layer C18 extraction disks (Empore) packed in 200μL pipette tip, which had been previously conditioned with 50μL100% methanol, 50μL 100% acetonitrile and 50μL 70% acetonitrile, 0.25% acetic acid. Filtered peptides were collected in a mass spectrometer vial and the peptides in the extraction disk were further eluted with 50μL 70% acetonitrile, 0.25% acetic acid and collected into the same mass spectrometry vial. Phosphopeptide-enriched samples were dried by vacuum centrifugation, solubilized in 3% acetonitrile, 4% formic acid, and one third of each sample was analyzed by LC-MS/MS.

### Liquid chromatography coupled to tandem mass spectrometry

Peptide samples were analyzed by nLC-MS/MS on a nanoAcquity UPLC (Waters) coupled to an Orbitrap Fusion Lumos Tribrid mass spectrometer (Thermo Fisher, San Jose, USA).Samples were loaded on a 100μm × 3-cm trap column packed with 3μm C18 beads (Dr. Maisch), separated on a 100μm × 30-cm capillary analytical column, packed with 1.9μm C18 beads (Dr. Maisch) and set at 50°C, using a 90-min reversed-phase gradient of acetonitrile in 0.125% formic acid, and online analyzed by mass spectrometry using data-dependent acquisition. Each cycle consisted of 3 sec where one full MS1 scan was acquired on the orbitrap at 120,000 resolution from 300 to 1575 m/z using an AGC of 7e5 and maximum injection time of 50ms followed by MS/MS dependent scans on most intense precursor m/z ions (only considering z = 2 to 5) until exhausting the 3sec cycle time, using 1.6 m/z isolation window, HCD fragmentation at 28 normalized collision energy, and acquired at 15,000 resolution on the orbitrap with an AGC of 5e4 (peptide samples) and AGC of 1e5 (phosphopeptide samples) with a maximum injection time of 22ms. Dynamic exclusion was enabled to exclude fragmented precursors from repeated MS/MS selection for 30sec. To increase coverage, phosphopeptide samples were injected twice, and the data from the two technical replicates were combined.

### Database searching, peptide quantification, phosphosite localization, and Rs calculation

MS data files for proteome samples were analyzed with MaxQuant^16^ (v.1.6.7.0) to obtain peptide identifications and quantifications, using the following parameters: protein sequence database *S.cerevisiae* downloaded from SGD in July 2014, LysC enzyme specificity (cleavage Ct to K), maximum of 2 missed cleavages, mass tolerance of 20ppm for MS1 and 20ppm for MS2, fixed modification of carbamidomethyl on cysteines, variable modifications of oxidation on methionines and acetylation on protein N-termini. Lysine residues were only allowed to be all light or all ^2^H_4_-Lys within the same peptide. Phosphoproteome samples were processed in MaxQuant similarly as above, with additional variable modification of phosphorylation on serine, threonine, and tyrosine residues. All searches were combined for MaxQuant filtering set to 1% FDR at the level of peptide spectral matches and protein.

Quantification values for heavy and light peptide features were extracted from the evidence.txt file. Quantification values for features corresponding to the same peptide sequence (e.g. same peptide identified at multiple charge states or fractions) were summed up. Phosphopeptide quantification features were aggregated to the phosphopeptide isoform level by summing features corresponding to the same peptide sequence (e.g. same peptide identified at multiple charge states or replicate injections) as well as overlapping peptide sequences sharing the same combination of modifications. For each phosphopeptide isoform we required the maximum localization probability to be greater than 75% for at least one site.

R_s_ values were calculated as log_2_ ratios of the quantification value for the temperature gradient treated divided by the respective quantification for the 30°C control. Peptide R_s_ distributions were median normalized to 0, and the same correction value derived for each replicate was applied to normalize the corresponding phosphopeptide isoform R_s_ distributions. Peptides and phosphopeptide isoforms with the 5% highest R_s_ standard deviation across replicates were excluded from the analysis. Protein R_s_ values were calculated as the median of peptide R_s_ for that protein, requiring a minimum of 2 peptides per protein, and each peptide observed in at least two replicates.

To identify phosphopeptide isoforms that have different R_s_ than their unmodified protein counterpart, we performed a t-test comparing phosphopeptide isoform R_s_ values (n=6) compared to protein R_s_ values (n=6) and assuming unequal variance. Phosphopeptide isoforms and protein counterparts were required to be observed in at least two replicates. P-values were corrected for multiple hypothesis testing using the Benjamini-Hochberg method^17^. All data analysis was conducted using R.

### Structure visualization and bioinformatics

Protein complex annotations were extracted from the CYC2008 resource^*18*^. Protein structure coordinates were downloaded from PDB and visualized and manipulated with PyMOL^19^. For PUP2 interface analysis, we extracted 20S proteasome protein structure from PDB 1RYP^20^. Protein interface structures for ARO8, TPI1, and GAPDH were extracted from PDB (4JE5^21^, 1NEY^22^, 3PYM^23^ respectively). To assess the stabilizing effect of S149 phosphorylation at the catalytic site of GAPDH, we aligned crystal structures of GAPDH with bound G3P (1NQO^24^) and inorganic phosphates (1GYP^25^) to a NAD-bound yeast GAPDH structure (3PYM^23^). Data on sequence conservation, protein interfaces, and predicted stability effects of mutations (ΔΔG_pred_) were obtained from the mutfunc resource^13^.

### Reanalysis of the Huang et al. data

Supplementary data from Huang et al.^2^ was used to calculate the correlation between T_m_ for phosphopeptides and proteins and learn about their statistical parameters. For data re-analysis, all MS files from the study were downloaded from MassIVE data repository (dataset identifier: MSV000083786), converted to open format mzXML files, and database searched with Comet^26^ (v.2018.01.4) to obtain peptide and phosphopeptide identifications, with the exception of “Bulk_6_2” which failed to convert. Database search parameters were: human protein sequence database from UniProt (UP000005640), mass tolerance of 50 ppm for precursor m/z and 0.2 Da for fragment ions, trypsin enzyme specificity (cleavage Ct to K, R, except for KP, RP), maximum of 2 missed cleavages, fixed modification of carbamidomethyl on cysteines and TMT10 (+229.1629) on lysines and peptide N-termini, and variable modifications of oxidation on methionines and acetylation on protein N-termini. Phosphorylation samples included variable modification of phosphorylation at serine, threonine, and tyrosine. Search results were filtered to a PSM 1% FDR with Percolator^27^. Phosphosite localization was conducted with an in-house C++ implementation of Ascore^28^ and sites with Ascore > 13 were considered confidently localized (P < 0.05). TMT10 reporter ion intensities were extracted from MS/MS scans using in-house TMT quantification software.

We attempted to replicate the analysis by Huang et al. by following the method description provided in their manuscript. Biological and technical replicates were treated equally. For each replicate, TMT reporter ion intensities for all peptide spectral matches from the proteome files were summed to the protein level, and TMT reporter ion intensities for phosphopeptide spectral matches were summed to the phosphopeptide isoform level. In addition, we used the same strategy to aggregate to peptide-level the TMT signals for PSMs mapping to the same unmodified peptide observed in the phosphopeptide-enriched samples. We implemented the TPP package in R to fit melting curves for proteins, phosphopeptide isoforms, and unmodified peptides in the phosphorylation-enriched sample. To recapitulate the reported results we had to conduct the fitting for all samples together (Supplementary Discussion). Melting curves were filtered for fitting R^2^ > 0.8. T-tests were conducted by comparing T_m_ values for phosphopeptide isoforms or unmodified peptides observed in the phosphopeptide-enriched samples to the unmodified protein T_m_ values, assuming equal variances and without multiple hypothesis correction (as implemented by Huang et al.). Of note, our reanalysis revealed that one of the phosphoproteome technical injections for biological replicate 5 was instead a repeated MS analysis of biological replicate 4.

### Data availability

The mass spectrometry proteomics data generated for this manuscript have been deposited to the ProteomeXchange Consortium via the PRIDE partner repository with the dataset identifier PXD016750. Reviewer username is; password 7aQ1gAbU.

## Supporting information

Supplementary Discussion

## ACKNOWLEDGEMENTS

We thank members of the Villén lab for scientific discussions, in particular Bianca Ruiz, Mario Leutert, and Alex Hogrebe. We thank Ariadna Llovet Soto and Jimmy Eng for software developments on the data analysis pipeline. I.R.S. and K.N.H were supported by NIH training grant T32HG000035. A.S.V. was supported by NIH training grant T32LM012419. Most of this work was supported by NIH grant R35GM119536 to J.V. The Villén lab is additionally supported by NIH grants R01AG056359, R01NS098329, and RM1 HG010461, Human Frontiers Science Program grant RGP0034/2018, a research program grant from the W.M. Keck Foundation, and the University of Washington Proteome Resource UWPR95794.

## AUTHOR CONTRIBUTIONS

I.R.S., K.N.H., R.A.R.-M, and J.V. conceived the study and designed the experiments. I.R.S. conducted the experiments with advice from K.N.H., R.A.R.-M. and J.V., and assistance from A.A.B. I.R.S. analyzed the data with advice from R.A.R.-M. and A.S.V. J.V. supervised the study. I.R.S. and J.V. wrote the paper and all authors edited it.

## COMPETING FINANCIAL INTERESTS

The authors declare no competing interests.

## SUPPLEMENTARY FIGURE LEGENDS

**Supplementary Figure 1.**
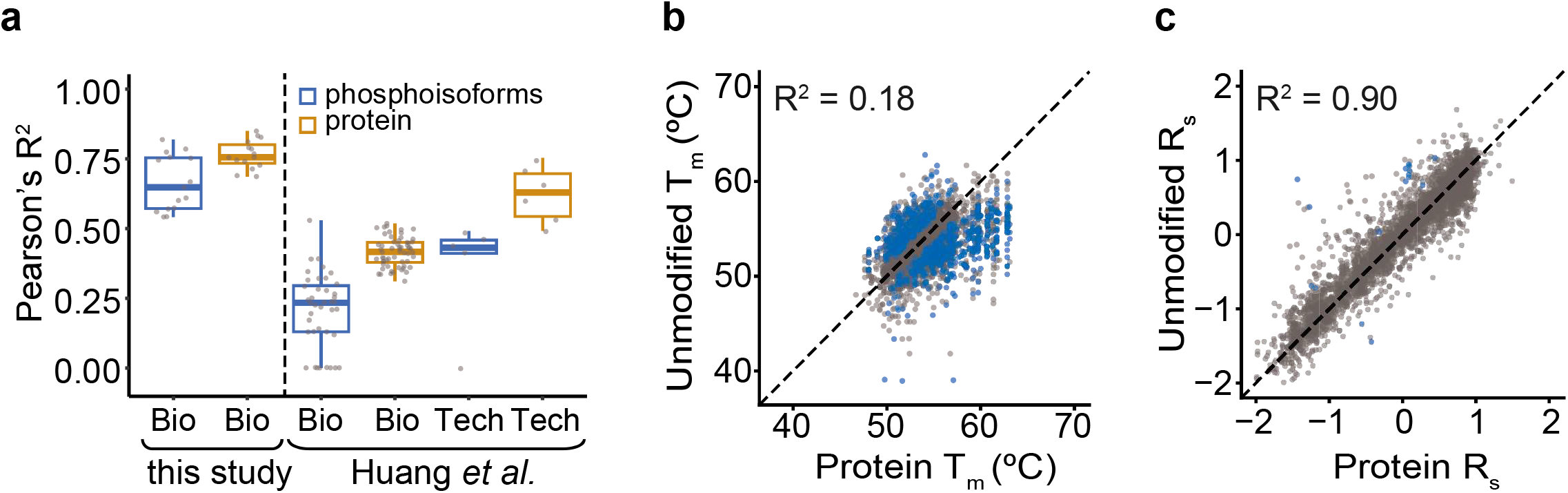
Reproducibility and robustness of Dali compared to HTP. **a**, Boxplot depicting pairwise Pearson correlations between biological and technical replicates for the HTP (T_m_ values) and Dali (R_s_ values) approaches. **b**, Scatter plot and Pearson correlation between the mean T_m_ for unmodified peptides observed in the phosphopeptide enriched samples (n=10) and the mean T_m_ for their corresponding proteins (n=11). Results from the Huang et al. data re-analysis conducted by us. **c**, Scatter plot and Pearson correlation as in (b) with R_s_ values obtained from the Dali method (n=6).

**Supplementary Figure 2:**
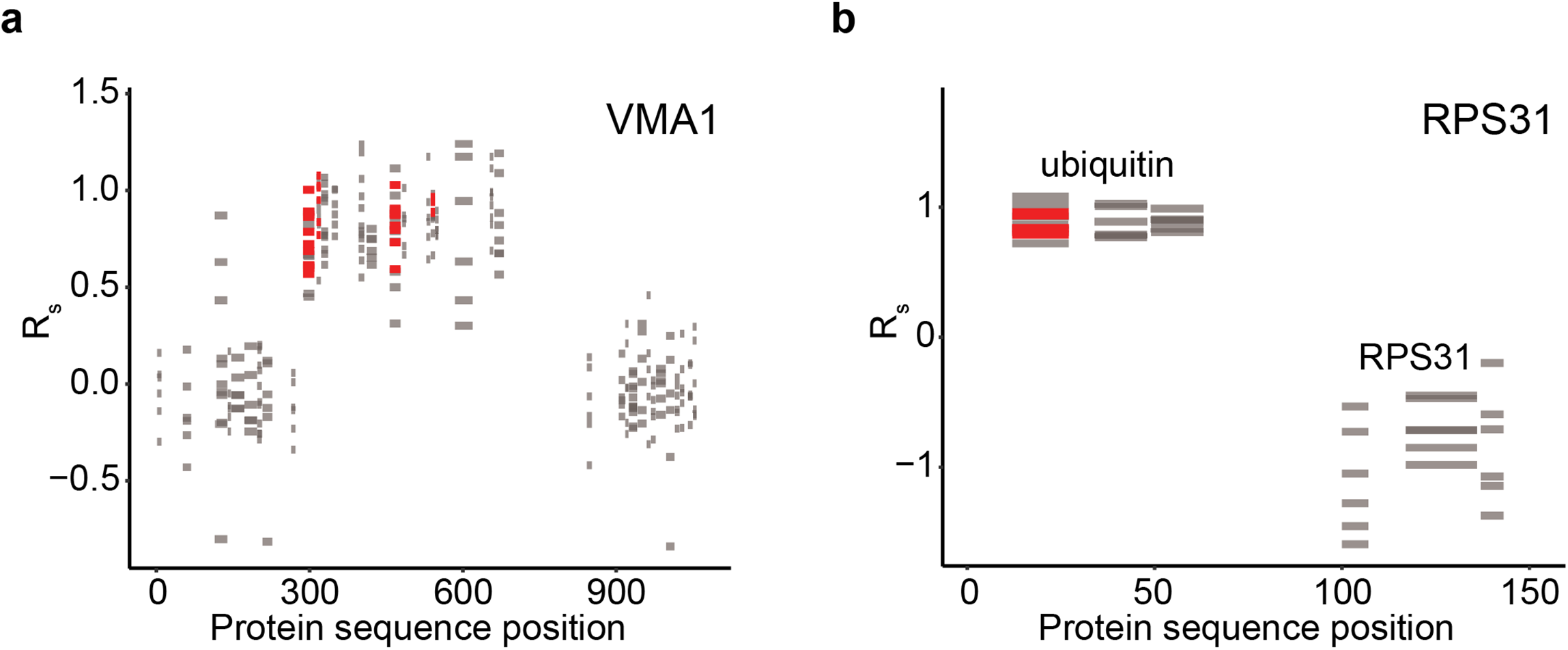
Examples of significant hits on proteins that undergo post-translational splicing or cleavage. **a**, R_s_ values for observed VMA1 unmodified peptides identified in phosphopeptide-enriched samples and proteome samples displayed across the length of VMA1. Spliced products from amino acid 2-283 and 738-1031 are joined to generate the V-type proton ATPase catalytic subunit A proteoform, extinguishing the 284-737 segment^7^. Peptides derived from the proteome samples are colored in gray and significant unmodified peptides found in the phosphopeptide-enriched sample are in red. **b**, Similar plot to (a) for RPS31, which is cleaved to generate ubiquitin (1-76 amino acid segment) and 40S ribosomal protein S31 (77-152 amino acid segment) proteins^6^.

**Supplementary Figure 3:**
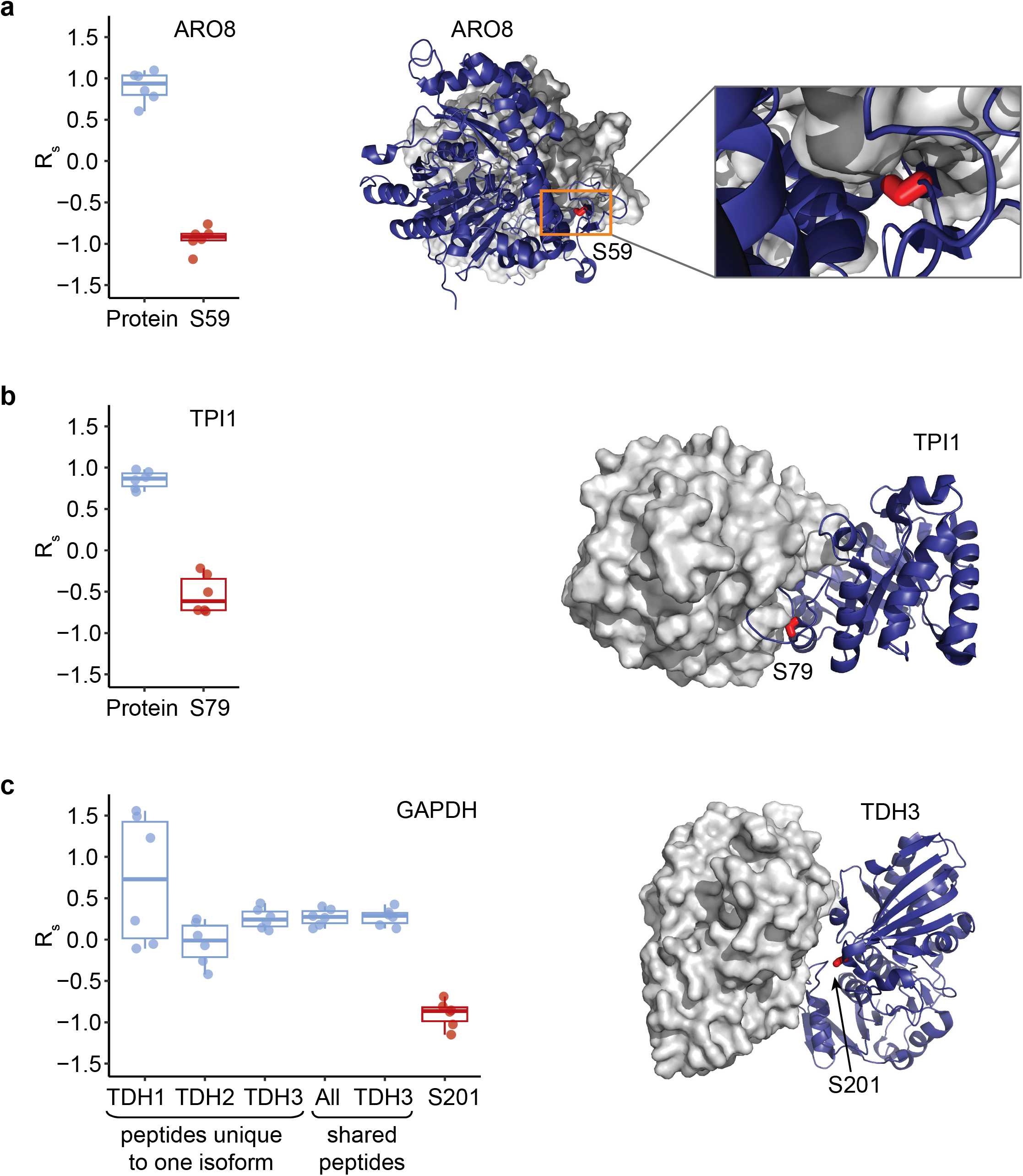
Examples of phosphosites that alter protein thermal stability and are located at protein interfaces. R_s_ boxplots for **a**, ARO8 S59, **b**, TPI1 S79, and **c**, GAPDH S201 phosphopeptide isoforms and their protein counterparts. ARO8 S59, TPI1 S79, and GAPDH S201 reside at dimerization interfaces as shown in the structures to the right (PDB accession: 4JE5, 1NEY, and 3PYM respectively). Phosphomimetic mutations ARO8 S59E and TPI1 S79E are predicted to disrupt protein interfaces (ΔΔG_pred_ = 3.78 and ΔΔG_pred_ =8.04 respectively). Additionally, TPI1 S79E mutation is predicted to alter protein conformational stability (ΔΔG_pred_ = 2.39). ΔΔG_pred_ > 2 is predicted to be destabilizing^12^.

**Supplementary Figure 4:**
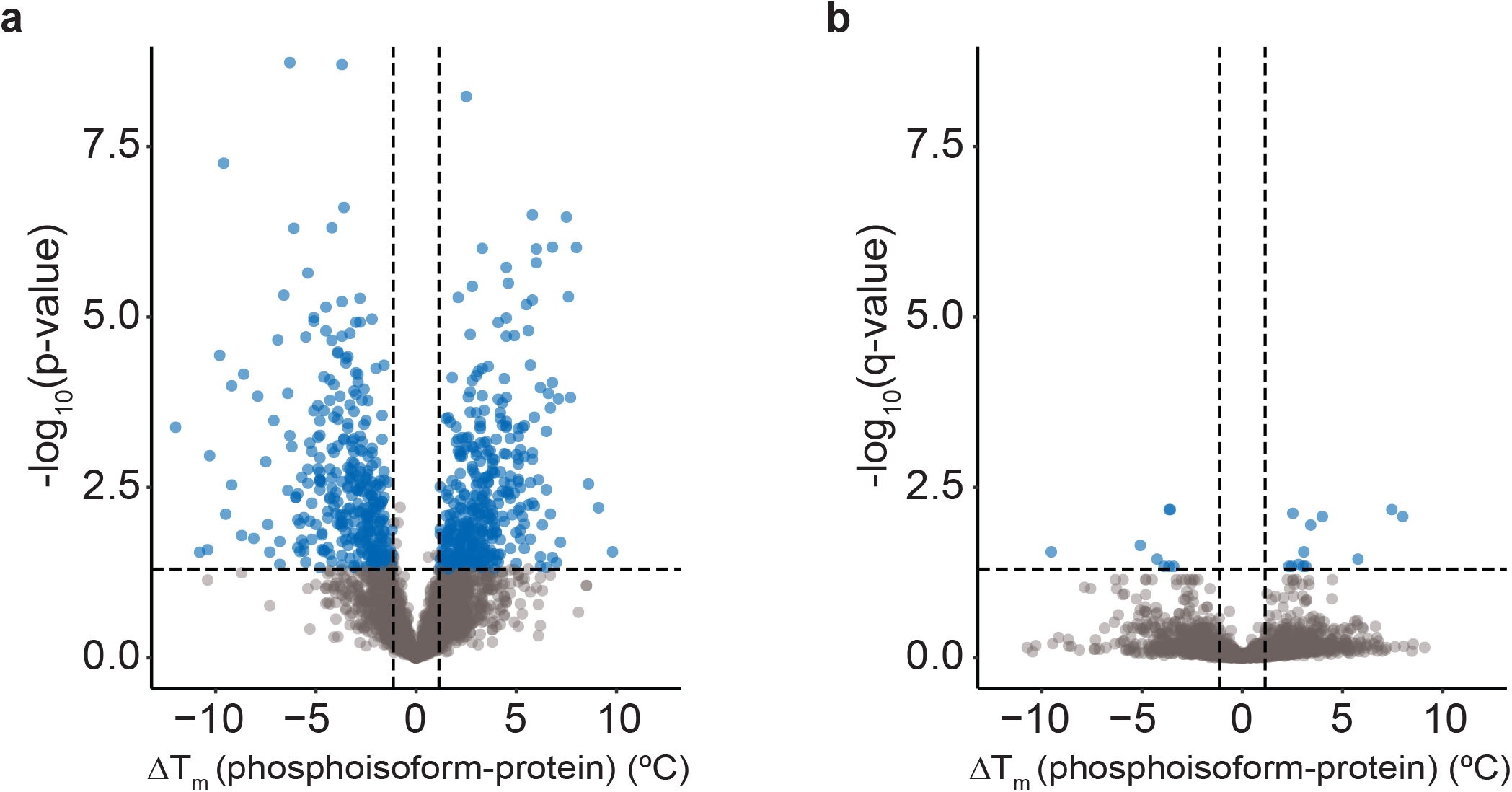
Phosphosites significantly altering protein thermal stability using two different statistical settings. Volcano plots showing ΔT_m_ for mean phosphopeptide isoform to mean protein counterpart in the x-axis, and the t-test probability in the y-axis. **a**, Huang et al. implementation shows a p-value because multiple hypothesis correction was not applied. Significant phosphopeptide isoforms (blue) are defined by p-value < 0.05. **b**, Our proposed analysis consolidates data from MS reanalysis prior to statistical testing, which is performed assuming unequal variances between phosphopeptide isoform and proteins and corrects p-values for multiple hypothesis testing. Significant phosphopeptide isoforms (blue) are defined by q-value < 0.05.

## Notes

http://proteomecentral.proteomexchange.org/cgi/GetDataset?ID=PXD016750

