## Supplementary Discussion for "Identification of phosphosites that alter protein thermal stability"

### SUPPLEMENTARY MATERIAL

#### SUPPLEMENTARY DISCUSSION

##### *Experimental workflow for phosphopeptide enrichment and TMT labeling*

HTP directly compares melting temperatures between phosphopeptides and their corresponding proteins. However, critical steps in the experimental workflow were conducted separately for phosphopeptides and proteins and also for the different temperature channels to trace melting curves, possibly introducing technical error. We indeed observe in the Huang et al. dataset low correlation between replicates (mean $R^2$ = 0.43 for protein, mean $R^2$ = 0.22 for phosphopeptides), between phosphopeptides and proteins ($R^2$ = 0.18 using the supplementary data provided; $R^2$ = 0.20 with our re-analysis), and between proteins and unmodified peptides identified in the phospho-enriched samples ($R^2$ = 0.18 with our re-analysis).

One potential source of error could be in the phosphopeptide enrichment, where previous work[1] has shown that the ratio of peptide to $TiO_2$ or IMAC stationary phase is an important parameter to optimize for successful enrichment, and deviations from the optimal ratio may result in variable peptide binding and/or recovery. In a thermal proteome profiling experiment, the soluble protein fraction recovered from the temperature treatment is very different across temperatures. Huang et al. enriched the derived peptides from each temperature separately but using the same amount of $TiO_2$ material. Our expectation with this setup would be highly variable phosphopeptide enrichment across the different temperatures. While we could not assess this directly, we observe variable phosphopeptide enrichment efficiencies across replicates ranging from 52 to 78%, with a substantial number of unmodified peptides in the phosphopeptide-enriched samples that appear to have significantly different $T_m$ compared to their reference protein.

Peptides for protein analysis and phosphopeptides were TMT-labeled and desalted separately. While we would expect uniform and high yield labeling reactions across all samples, the workflow does not control for any potential differences. To minimize all these sources of error, we suggest conducting TMT labeling of peptides first and apply phosphopeptide enrichment after the samples from the different temperature channels have been mixed. This scheme would minimize the distortion of the melting curves for phosphopeptides[2].

Our workflow introduces the isotopically-labeled 30$^o$C sample to control for any technical differences occurring after temperature treatment, including phosphopeptide enrichment and sample cleanup. As a result, we observe high correlation between replicates, between proteins and unmodified peptides identified in the phosphorylation enriched sample ($R^2$=0.90), and between phosphorylation isoforms and proteins ($R^2$= 0.79).

##### *TMT ratio and $T_m$ compression*

HTP applies MS2-based TMT quantification of unmodified proteome and phosphoproteome samples in single shot injections. Many studies[3,4] have reported TMT ratio compression, and attributed this to interfering TMT signals derived from peptides that have been co-isolated for MS/MS fragmentation with the precursor of interest. We expect interference and ratio compression would increase along with sample complexity, and would be high for the experimental settings used in the Huang et al. study, consisting of whole human proteome measurements over 2h (replicates 1 and 2) or 4h (replicates 3-6) LC-MS/MS run times. Indeed, we observe the technical replicates of the longer runs to be better correlated ($R^2$=0.7) than those for the shorter runs ($R^2$=0.5). We expect TMT ratio compression will translate into a compression of protein melting curves and melting temperatures trending towards the sample $T_m$ average. The low reproducibility between LC-MS/MS repeat injections (average proteome $R^2$=0.62, average phosphoproteome $R^2$=0.36 , Supplementary Fig. 1a) supports that TMT ratio compression could be a source of error.

Several approaches have been described to mitigate TMT ratio compression including the use of an orthogonal fractionation of the proteome to reduce complexity of each MS injection, filtering the data to only include TMT quantifications derived from isolating a single predominant precursor signal[3], decreasing the isolation window, and/or applying a sequential isolation step in the MS to quantify TMT reporters in an MS3 scan[5].

In our method, we fractionated the proteome into 5 fractions to reduce complexity and obtain deeper coverage. We also chose to use a SILAC-based quantification approach, which has proven to provide more accurate phosphopeptide quantifications than TMT[6] and does not suffer from ratio compression. We think quantification accuracy becomes more important with peptide or phosphopeptide isoform analysis than with protein analysis, given that the final quantitative measurement aggregates only one to a few measurements.

##### *TPP melting curve fitting*

We found that in order to recapitulate the Huang et al result of the phosphopeptide $T_m$ and protein $T_m$ having the same median $T_m$ all data must be fit together, so sample-to-sample normalization in the TPP package[7] could adjust all phosphorylation and unmodified protein sample consensus curves towards a single representative curve. We believe that this is not the correct way to apply the TPP package because it may obviate global differences in protein and phosphoprotein melting. Separate analyses of proteins and phosphopeptide isoforms for the Huang et al. data indeed revealed existing differences in the average melting curve. Rather, unmodified peptides that are common between enriched and unenriched samples could be used for data normalization[2].

##### *Statistical analysis*

The study by Huang et al. encompassed five biological replicates for phosphorylation analysis and six biological replicates for unmodified proteome analysis, and each was analyzed twice on the mass spectrometer. When conducting the t-test to compare the $T_m$ of phosphopeptide isoforms and proteins, the authors treated mass spectrometer repeat injections as independent biological replicates, thus artificially increasing the power of their statistical tests. Additionally, the authors led with the assumption that the $T_m$ variances of the unmodified protein and the phosphoprotein were equal, while we calculated these as 2.4 and 9.5 respectively. Lastly, the authors did not adjust p-values for multiple testing.

With these considerations, we reimplemented the t-test on the Huang et al. dataset by applying median consolidation of reanalyzed samples, assuming unequal variances, and adjusting p-values for multiple testing using the Benjamini-Hochberg method. This resulted in a dramatic decrease in the number of phosphopeptide isoforms that significantly alter protein $T_m$, from 719 to 20 (Supplementary Fig.4).

##### *Limitations of the SILAC-based method*

We acknowledge some limitations inherent to the approach we use. First, each sample combines two independent cell cultures grown in light and heavy SILAC media. It is possible that there may be some differences in protein abundance that may introduce variability across replicate measurements of protein and phosphopeptide isoform $R_s$ values. We partially control for this by swapping the isotopic labeling scheme for half of the replicates and partially mitigate the issue by removing the 5% most variable data.

A second limitation is on the magnitude of the $R_s$ values, which provide a measure of the relative stability of the protein (or phosphoprotein) to unfold and aggregate under a temperature gradient centered around 50$^o$C *vs.* 30$^o$C. Therefore, $R_s$ values are relative to the $T_m$ of each protein, and our ability to detect changes to $R_s$ values depends on the temperature gradient chosen, relative to the protein $T_m$. In our study, we selected a $T_m$ gradient centered around the median $T_m$ for the *S. cerevisiae* proteome. Thus, we expect to have missed in our study functionally relevant phosphosites in proteins with extreme melting temperatures. In order to capture these, additional experiments would be required where the temperature gradient is globally shifted towards lower or higher temperatures.
